# Mechanosensory input during circuit formation shapes Drosophila motor behavior through Patterned Spontaneous Network Activity

**DOI:** 10.1101/2021.03.21.436277

**Authors:** Arnaldo Carreira-Rosario, Ryan A. York, Minseung Choi, Chris Q. Doe, Thomas R. Clandinin

## Abstract

Neural activity sculpts circuit wiring in many animals. In vertebrates, patterned spontaneous network activity (PaSNA) generates sensory maps and establishes local circuits ^1–3^. However, it remains unclear how PaSNA might shape neuronal circuits and behavior in invertebrates. Previous work in the developing *Drosophila* embryo discovered spontaneous muscle activity that did not require synaptic transmission, and hence was myogenic, preceding PaSNA ^4–6^. These studies, however, monitored muscle movement, not neural activity, and were therefore unable to observe how myogenic activity might relate to subsequent neural network engagement. Here we use calcium imaging to directly record neural activity and characterize the emergence of PaSNA. We demonstrate that the spatiotemporal properties of PaSNA are highly stereotyped across embryos, arguing for genetic programming. Consistent with previous observations, we observe neural activity well before it becomes patterned, initially emerging during the myogenic stage. Remarkably, inhibition of mechanosensory input as well as inhibition of muscle contractions results in premature and excessive PaSNA, demonstrating that muscle movement serves as a brake on this process. Finally, using an optogenetic strategy to selectively disrupt mechanosensory inputs during PaSNA, followed by quantitative modeling of larval behavior, we demonstrate that mechanosensory modulation during development is required for proper larval foraging. This work thus provides a foundation for using the *Drosophila* embryo to study the role of PaSNA in circuit formation, provides mechanistic insight into how PaSNA is entrained by motor activity, and demonstrates that spontaneous network activity is essential for locomotor behavior. These studies argue that sensory feedback during the earliest stages of circuit formation can sculpt locomotor behaviors through innate motor learning.

**Highlights:**

- PaSNA in the *Drosophila* embryonic CNS is spatiotemporally stereotyped
- Mechanosensory neurons negatively modulate PaSNA
- Embryonic PaSNA is required for larval locomotor behavior

## Results

### PaSNA in the Drosophila embryo

Motor movements begin in the embryo as uncoordinated twitching at stage 16, followed by larger scale movements that progressively become stronger and more organized prior to hatching approximately 7 hours later (Figure 1A). To characterize the emergence of neural activity across these stages, as well as to make comparisons between animals, and to facilitate rapid screening of neural and molecular perturbations, we developed a wide-field imaging preparation in which we could monitor neural activity in 25-35 embryos simultaneously (Figure 1B; Methods). We expressed the genetically encoded calcium indicator GCaMP6s in all neurons, while co-expressing nuclear TdTomato to allow for ratiometric imaging, and acquired images every 7 seconds from the myogenic stage through hatching (Video S1). Under these imaging conditions, 95% of control animals hatched (n = 60), demonstrating that this preparation does not disrupt normal development. Finally, to correct for small variations in the developmental timing of individual embryos, we monitored ventral nerve cord (VNC) condensation and normalized developmental stage by computing the ratio of the length of the embryo to the length of the central nervous system (CNS) (Figure S1A), following standard methods ^7,8^.

**Figure 1.**
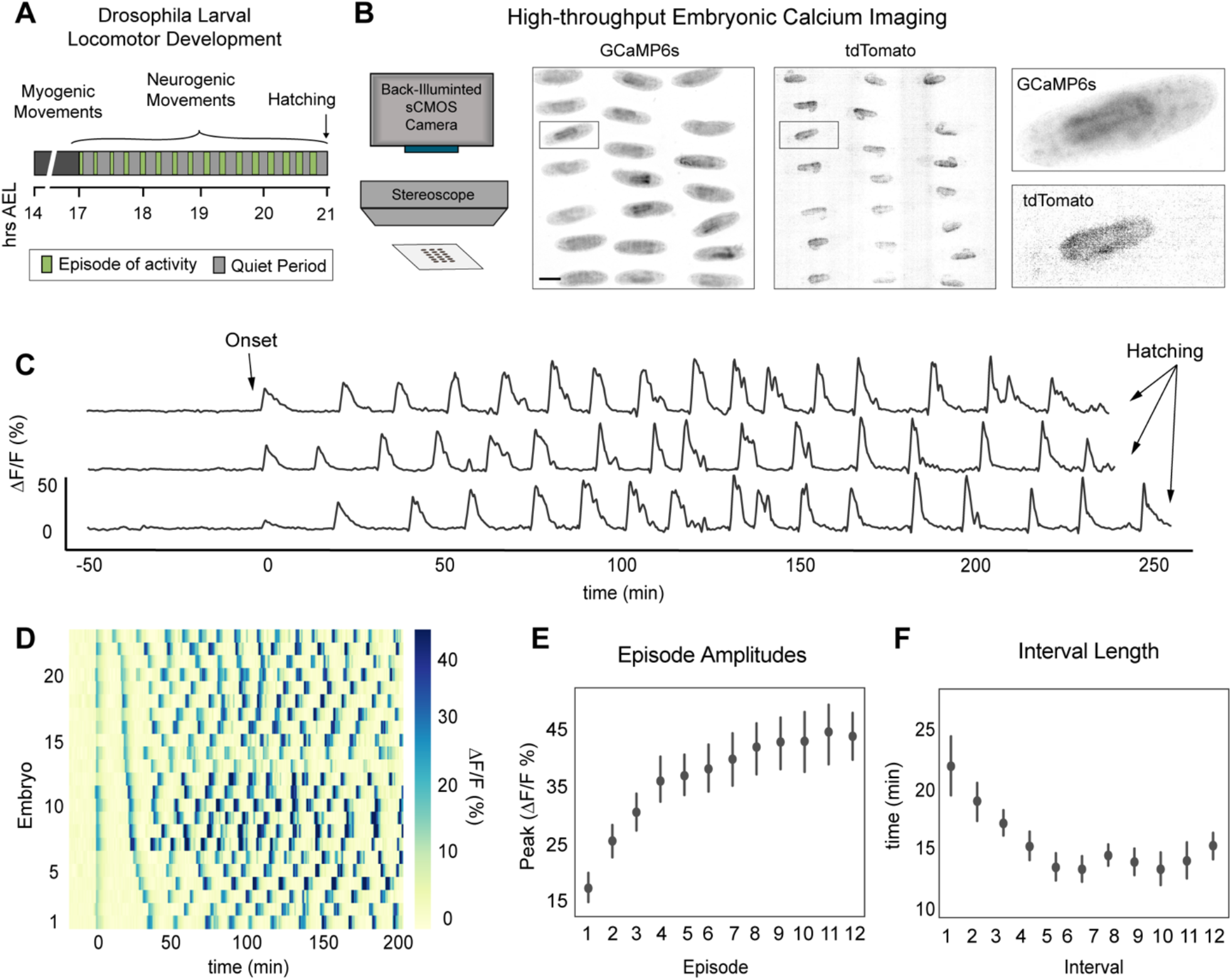
Characterization of patterned spontaneous network activity in Drosophila embryo. **(A)** Schematic of *Drosophila* larval locomotor development. Time in hours after egg laying (hrs AEL). **(B)** Schematic of high-throughput imaging system (left). Images of GCaMP6s and tdTomato signal across the imaging field, color inverted for visualization (right). Scale bar: 200 μm. **(C)** GCaMP6s:TdTomato ΔF/ F traces from three individual embryos. **(D)** Raster plot for PaSNA trimmed at 200 minutes post-onset, sorted by distance between first and second peak, with each trace corresponding to an individual embryo. Increasingly strong movements prevent accurate measurements at later stages. ΔF/ F heat map scale to the right. **(E)** ΔF/ F peaks for episodes 1 through 12 (n = 23). **(F)** Quantification of the first eleven interbout interval lengths (n = 23). For (**E**) and (**F**), points represent mean and lines depict the 95% confidence interval (CI). For genotype information see Table S1.

Consistent with the pattern of muscle movements ^5^, we observed episodes in which intracellular calcium concentrations increased in many neurons and their processes (Figure 1C, D; Video S1). Strikingly, the timing of the first large wave of neural activity was highly consistent from animal to animal, appearing at a length ratio of 2.2 (95%CI [0.06, 0.06]) (Figure S1B), corresponding to early stage 17. Aligning calcium traces by the timing of the first episode revealed that the overall PaSNA pattern was qualitatively and quantitatively similar across all embryos (Figure 1D-F). In particular, a total of 17 PaSNA episodes (95%CI [0.99, 0.99]) that occurred over 275 minutes (95%CI [18.3, 18.3]) preceded hatching (Figure S1B, C). Moreover, the size and duration of each wave of activity consistently increased over the first eight waves, before stabilizing (Figure 1E). Finally, in parallel with the increasing strength of the early episodes, the interbout interval dramatically decreased over the first five episodes of PaSNA from 21.8 minutes (95% CI [2.3,2.5]) to 13.3 minutes (95% CI [1.1,1.2]) (Figure 1 F). Our stereotypy analysis revealed that the observed interbout intervals were significantly more stereotyped across embryos than random (Figure S1D). Taken together, these data show that PaSNA is highly stereotyped from embryo to embryo, suggesting that PaSNA is genetically encoded.

### Spatiotemporal properties of the initial PaSNA episode

We focused next on the first episode of PaSNA, a period of particular interest given that it represents the transition from myogenic to neurogenic movement. To what extent is the pattern of neural activity underlying the first episode stereotyped across embryos? To investigate the spatiotemporal patterns of neural activity during single episodes of PaSNA, we developed a two-photon (2P) microscopy preparation to image embryos expressing pan-neuronal GCaMP6s and TdTomato. This system allows for imaging of the entire VNC for two hours at cellular resolution, acquiring imaging volumes at 2.6 Hz. Embryos survive imaging, hatch and become adult flies (n = 8). To unequivocally identify the first episode of PaSNA, we began imaging at least 30 minutes before the neurogenic phase. Preceding the first episode of PaSNA, we observed sporadic neuronal firing throughout the VNC, an activity pattern we refer to as flickering. This activity was observed during the 30 minutes before the first PaSNA episode, thus appearing during the myogenic phase of movement ^5^. After this, the first PaSNA episode began, and comprised three phases, namely *localized initiation, propagation* and *peak activity* (Figure 2A; Video S2). During the first phase, we observed increased levels of neural activity within a stereotyped region, marking the *localized initiation* of PaSNA and defining the start of neurogenic activity. During the second phase, we observed a single wave of neural activity that traversed the VNC and defined *propagation*. During the third phase, we observed a period of *peak activity* along the VNC that persisted for approximately 80 seconds. Activity then returned to basal levels where an interbout interval containing flickering activity persisted until the next episode.

**Figure 2.**
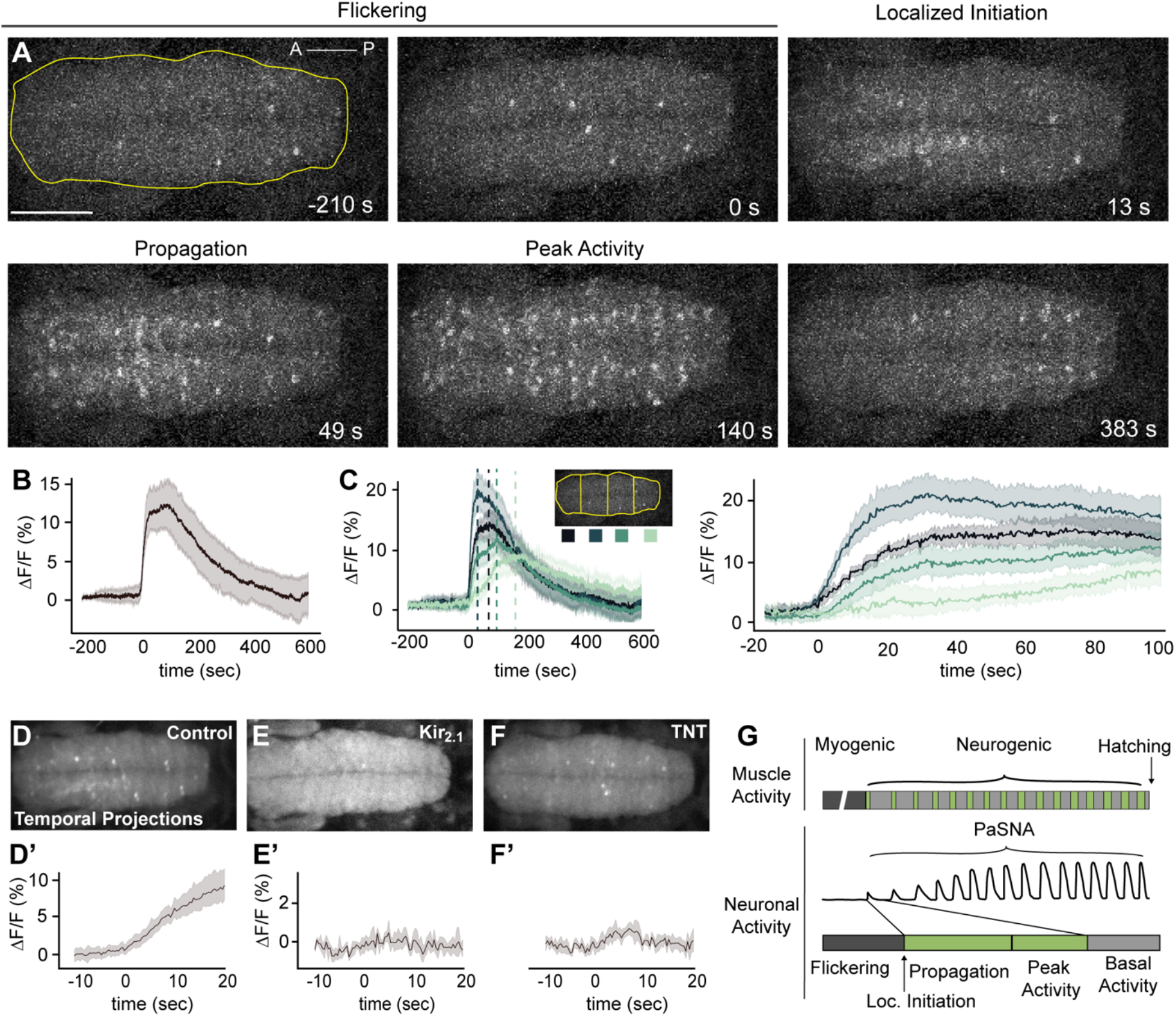
Spatiotemporal and network properties of a single PaSNA episode. **(A)** Image frames during the first episode of PaSNA in an embryo. Stages labeled on top. Images are maximum intensity projections from an embryonic VNC expressing pan-neuronal GCaMP6s. Time stamps are relative to the positive inflection point caused by the activity burst. Yellow line delineates the VNC, with the ROI used for Panel B. Scale bar: 50 μm. **(B)** ΔF/F trace of the entire VNC during the first episode of PaSNA (n = 8). **(C)** ΔF/F of the color-coded four ROIs. Left displays −200 seconds to 600 seconds; right displays from −20 to 100 seconds relative to the initiation of PaSNA. **(D-F’)** Temporal projections (top) and ΔF/F VNC traces (bottom) for 30 seconds near the localized initiation time of the episode for control embryos (n = 8) (**D**), embryos expressing Kir_2.1_ pan-neuronally (**E**) (n = 5) and embryos expressing TNT pan-neuronally (F) (n = 6). **(G)** Schematic of *Drosophila* larval locomotor development showing activity at the muscle (top) and neuronal level (bottom). For all time series, dark lines represent the mean, while shading depicts the 95%CI. For genotype information see Table S1.

The focal activity observed during the *localized initiation* phase prompted us to examine whether the location of this event was invariant across embryos. Analysis of neural activity within ROIs along the anterior-posterior (A-P) axis of the VNC showed that PaSNA always initiated in the anterior region of the VNC (Figure 2C) (n =8). Furthermore, in 100% of embryos, activity initiated in one of the two most anterior ROIs, a region spanning the thoracic segments. After initiation, activity always propagated along the A-P axis. Strikingly, the wave of neural activity propagated slowly, reaching the most posterior region of the embryo approximately 75 seconds after localized initiation, corresponding to a propagation speed of less than 2 μm per second. Lastly, in all embryos, the more posterior regions were the last to return to basal, flickering activity. Together, these observations demonstrate that the initial episode of PaSNA is spatiotemporally patterned.

### The role of neural activity in initiating PaSNA

Next, we examined the role of neural activity in the initiation of PaSNA. To test whether neuronal depolarization caused the observed calcium transients recorded with GCaMP, we inhibited depolarization in all neurons through pan-neuronal expression of the inward-rectifier potassium channel Kir_2.1_ ^9^. As expected, this abolished flickering during the myogenic phase, as well as all three phases of PaSNA, indicating that PaSNA is a voltage-dependent process (Figure 2E; Video S3). Next, we tested whether PaSNA is driven by chemical synapses by inhibiting synaptic transmission using tetanus toxin (TNT) ^9^. Pan-neuronal expression of TNT had no effect on flickering during the myogenic phase, but prevented all three phases of PaSNA including the propagating waves of neural activity (Figure 2F; Video S4). This demonstrates that the flickering preceding PaSNA emerges in the absence of synaptic transmission, and thus is likely due to the intrinsic membrane excitability of individual neurons. Together, these results show that while neuronal depolarization and chemical synaptic transmission are both crucial for PaSNA, only depolarization is required for flickering.

### Mechanosensory input negatively modulates PaSNA

We next sought to determine whether the initial, myogenic phase of spontaneous muscle movement might be functionally coupled to the initiation of PaSNA. We reasoned that this coupling could occur through sensory feedback via proprioceptors. Therefore, we first asked which proprioceptive neurons are active during the myogenic phase. We used the calcium integrator system CaLexA, a method for transcriptionally labeling active neurons (Figure 3A) ^10^. To restrict this system to only those neurons that are active in the absence of synaptic transmission, we used pan-neuronal expression of TNT (Figure 2F; Figure 3A). Thus, the CaLexA reporter can only be induced in neurons that are active during the myogenic phase, allowing selective labeling of single cells (Figure 3A). Strikingly, every embryo displayed high levels of CaLexA expression in mechanosensory chordotonal (mechano-ch) neurons in most segments (n = 30) (Figure 3B-E). Specifically, observed expression in lch5 in every hemisegment as well as expression in lch1 and vchA/B in some segments. Notably, none of these embryos showed CaLexA signal in any other proprioceptive neurons.

**Figure 3.**
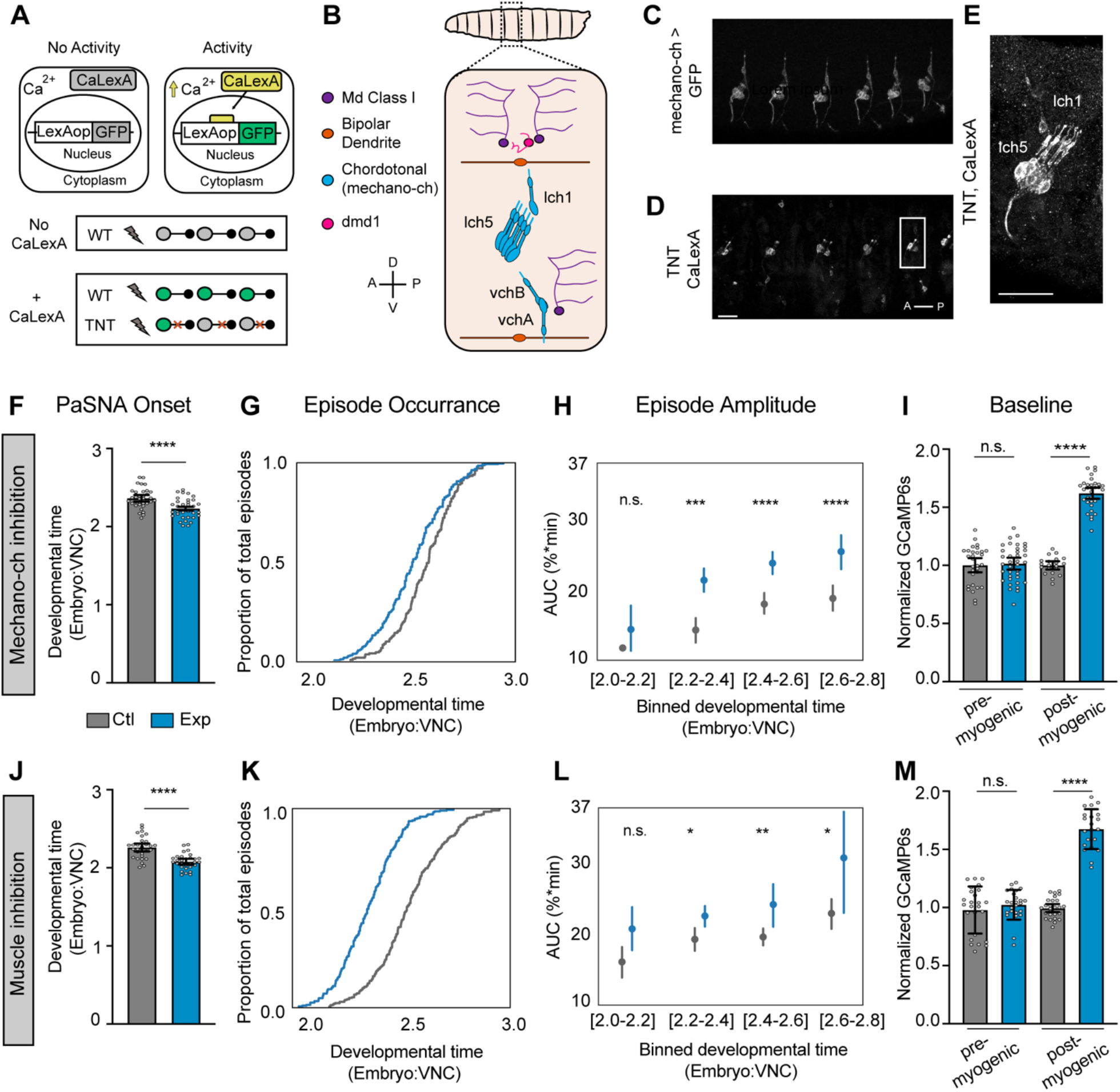
Mechanosensory neurons modulate the amplitude of PaSNA episodes. **(A)** Schematic illustration of the experiment using CaLexA to reveal neural activity during the myogenic phase. **(B)** Schematic of an embryonic anterior body wall hemisegment showing all proprioceptive neurons. There are eight mechanosensory chordotonal neurons (mechano-ch [blue]). Five of these form a laterally located cluster (lch5). A solitary mechano-ch is located dorsal to lch5 (lch1), and a pair of mechano-ch neurons is located ventrally (vchB and vchA). Anatomical coordinates: anterior (A), posterior (P), dorsal (D) and ventral (V). **(C)** Expression of the mechano-ch driver *inactive* (*iav*) along several body wall segments. **(D-E)** CaLexA driving GFP expression in a 19 hrs AEL embryo expressing pan-neuronal TNT (n= 30 embryos). Note expression in lch5 in every hemisegment as well as expression in lch1 and vchA/B in some segments. Scale bars: 20μm. **(F-I)** Measurements of the timing and intensity of PaSNA in control embryos (gray) and experimental embryos expressing TNT in mechano-ch neurons (blue). **(F)** Quantification of PaSNA onset (n = 36 control; n = 33 experimental). **(G)** Cumulative occurrence of the first twelve episodes plotted as the proportion of total episodes across developmental time (n = 17 control; n = 32 experimental). **(H)** Area under the peak curve (AUC) quantification for the first twelve episodes plotted against developmental time. Values were binned based on developmental time (n = 17 control; n = 32 experimental). **(I)** Quantification of GCaMP6s baseline levels normalized against control mean before (14hrs AEL; n = 30 control; n = 37 experimental) and after (10 minutes before PaSNA onset; n = 20 control; n= 32 experimental) the myogenic phase. **(J-M)** Measurements of the timing and intensity of PaSNA in control embryos (gray) and experimental embryos expressing Kir_2.1_ in muscles (blue). **(J)** Quantification of PaSNA onset (n = 36 control; n = 30 experimental). **(K)** Cumulative occurrence of the first twelve episodes plotted as the proportion of total episodes across developmental time (n = 28 control; n = 28 experimental). **(L)** AUC quantification for the first twelve episodes plotted against binned developmental time (n = 28, control; n = 28 experimental). **(M)** Quantification of GCaMP6s baseline levels normalized against control mean before (n = 25 control; n = 25 experimental) and after (n = 26 control; n = 21 experimental) the myogenic phase. For (**H**) and (**L**), points represent mean and lines depict the 95% confidence interval. For all bar graphs the mean and 95% CI are displayed. ****p<0.0001, ***p<0.001, **p<0.005, *p<0.05. For (**F**) and (**J**) we used two-sample t-tests. For (**H**) and (**L**) we used two-sample t-tests with Holm-Bonferroni correction. For (**I**) and (**M**) we used two-sample Welch’s t-tests to account for differences in variance. For genotype information see Table S1.

Mechano-ch neurons detect muscle stretch, relaying proprioceptive signals used to regulate larval crawling speed ^11–14^. Additionally, these neurons have been previously linked to embryonic neural circuit formation ^12^, making them ideal candidates for coupling muscle movements to PaSNA. To test this idea, we first silenced mechano-ch neurons and examined whether this perturbation altered the timing or amplitude of PaSNA episodes. To do this, we used the *inactive* (*iav*) enhancer to express TNT in mechano-ch neurons, and monitored neural activity throughout PaSNA using wide-field imaging of pan-neuronally expressed GCaMP6s. Consistent with previous work demonstrating that blocking all sensory neuron function did not prevent the emergence of muscle movements^5^, PaSNA was not abolished after mechano-ch silencing. Strikingly, however, PaSNA started prematurely in these embryos (Figure 3F). This led to embryos experiencing more episodes of PaSNA earlier in development (Figure 3G), as well as increasing PaSNA duration, and the total number of episodes (Figure S2A,B). Importantly, the amplitude of most PaSNA episodes was increased in these embryos as compared to controls (Figure 3H). Lastly, interbout intervals remain largely unchanged (Figure S2C).

We reasoned that if mechano-ch neurons were coupling muscle contraction to PaSNA, there might be an effect of blocking the synaptic transmission in these cells during the myogenic phase. To test this, we examined the baseline fluorescence of GCaMP6s (relative to nuclear TdTomato expressed on the same RNA transcript), as a proxy for the intracellular calcium concentration and membrane excitability, before and after the myogenic phase. As expected, inhibiting mechano-ch neurons had no effect on baseline fluorescence when muscles are yet to contract, before the myogenic phase (Figure 3I). Strikingly, inhibiting mechano-ch neurons dramatically increased the baseline fluorescence signal of GCaMP6s after muscles have contracted during the myogenic phase (but before PaSNA began; Figure 3I). Such an increase in baseline GCaMP6s signal is consistent with higher intracellular calcium concentrations and increased membrane excitability, providing a potential explanation for the premature onset and increased amplitude of PaSNA seen when mechano-ch neurons are inhibited. We note that this change in baseline GCaMP6s fluorescence cannot be accounted by changes in protein expression, as these measures are normalized relative to TdTomato in every cell. Thus, mechano-ch neurons act during the myogenic phase. Finally, to complement these results, we repeated these experiments in embryos expressing the inward rectifying channel Kirinterspersed by low-activity periods _2.1_ in mechano-ch neurons. However, while we observed significant increases in the amplitudes of initial PaSNA episodes (Figure S2G), overall effects were modest, suggesting that this functional inhibition was incomplete, and further obscured by genetic background effects that increased baseline GCaMP6s fluorescence (Figure S2L).

If mechano-ch neurons are coupling muscle movements to PaSNA, inhibiting muscle contractions should have similar effects to inhibiting mechano-ch neurons. To test this, we inhibited muscle contraction by expressing Kir_2.1_ using a muscle specific driver ^15^, while expressing GCaMP6s pan-neuronally. Strikingly, preventing muscle contraction caused premature PaSNA onset, increased the amplitude of PaSNA episodes, and led to higher baseline fluorescence in post-myogenic but not pre-myogenic embryos (Figure 3J-M). Thus silencing muscle contraction, or proprioceptive neuron function lead to premature onset of PaSNA and increased amplitude of PaSNA episodes, strongly suggesting that muscle contractions induce mechano-ch activity during the myogenic phase to negatively modulate PaSNA.

### Developmental inhibition of mechanosensory input leads to abnormal larval behavior

Our observation that silencing mechano-ch neurons increased PaSNA raised the question of whether this change in PaSNA had behavioral consequences, and more specifically, whether these transient changes in PaSNA resulted in long-term behavioral deficits. We inhibited mechano-ch activity transiently (from the late myogenic phase through to the end of PaSNA) by using the iav enhancer to express the *G. theta* anion channelrhodopsin 1 (GtACR1) ^16^, and examined larval behavior 24hrs after hatching using the Frustrated Total Internal Reflection-based imaging method (FIM; Figure 4A)^17^. We employed Time REsolved BehavioraL Embedding (TREBLE) to characterize potential behavioral differences between the optogenetically silenced condition and the control. TREBLE is a quantitative framework for identifying structure in behavior by collecting features (such as larval posture or velocity) into temporal windows and embedding these into a low-dimensional space (Figure 4B). As previously shown^18^, we found that the major components of the larval foraging ethogram^19,20^ can be captured in a 2-dimensional space using TREBLE (Figure 4B, C). In this 2-dimensional space, crawling is represented by an oscillator with directional movement (Figure S3) and is connected to regions corresponding to pausing and turning (Figure 4C).

**Figure 4.**
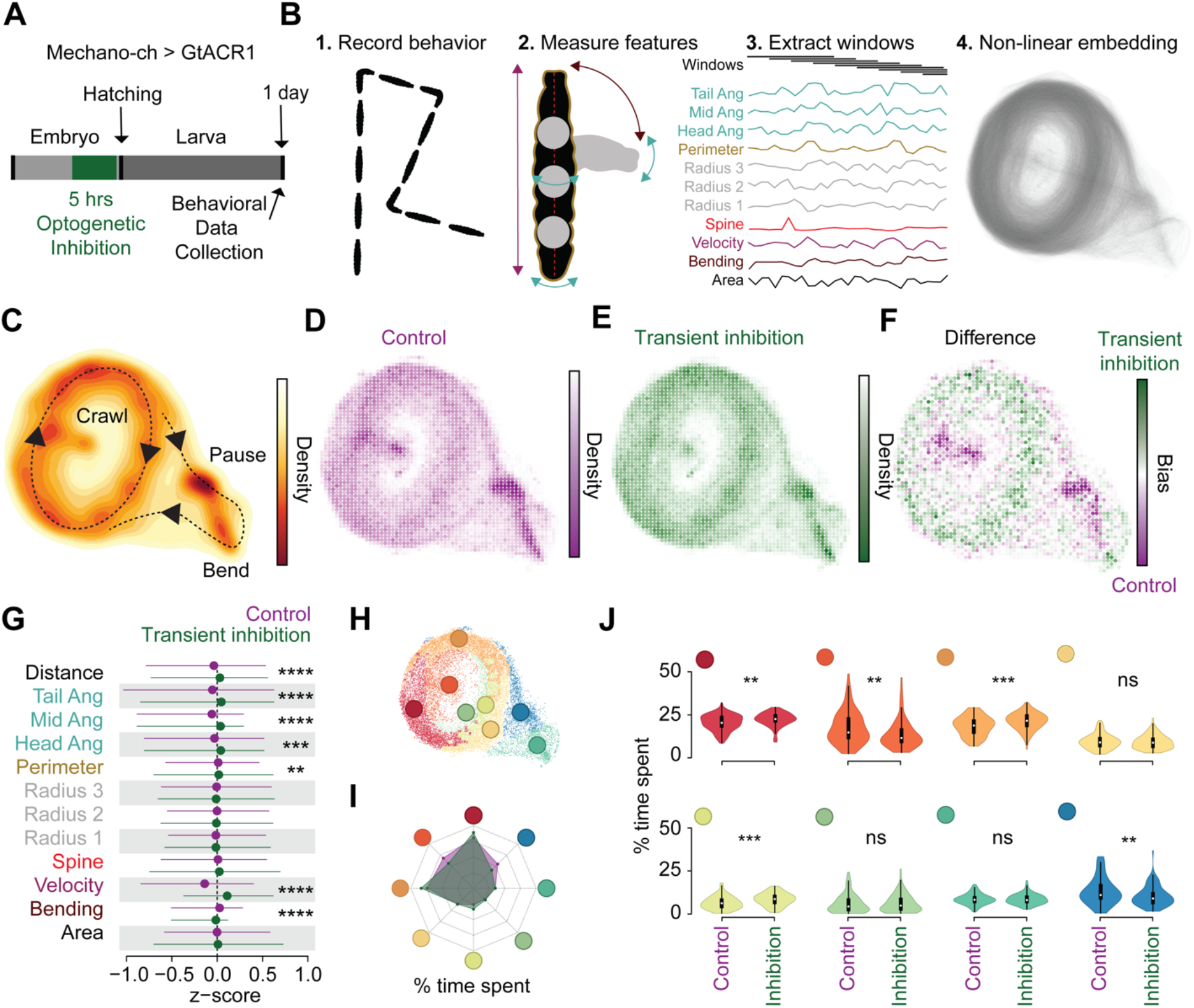
Temporal embryonic inhibition of mechanosensory input leads to abnormal larval behavior. **(A)** Schematic of experimental design. **(B)** Workflow for Time Resolved BehavioraL Embedding (TREBLE) (Methods). **(C)** Probability density function of larval locomotor space plotted as a heatmap. Behaviors annotated qualitatively. Density scale to the right. **(D-E)** Bin-wise occurrence distributions for control (n= 84) (D) and transient inhibition (n = 97) (E) groups. **(F)** Difference map between control (purple) and transiently inhibited (green) animals. Bias scale to the right. **(G)** Comparison of primary behavioral features between control (purple) and transiently inhibited (green) larvae. **(H)** Behavioral space colored via Louvain clusters (Methods). **(I)** Radar chart comparing the percentage of time spent in each of the Louvain clusters for control (purple shade) and transient inhibition (green shade) groups. **(J)** Differences in occurrence in each behavioral cluster between control and transiently inhibited animals. **** p < 0.0001, *** p< 0.001, ** p < 0.01. For (**G**) we used trial-wise Kruskal-Wallis test with Bonferroni correction. For (**J**) we used trial-wise Kruskal-Wallis test. For genotype information see Table S1.

In the TREBLE approach, both control and transiently inhibited larvae were used to generate a single, common behavior space (n = 181 total; 84 control larvae, 97 transiently inhibited larvae); 179,409 windows; see Methods) where we could directly infer behavioral variation via differences in the likelihood that either control or experimental larvae occupied specific regions of the space (Figures 4D-E). Control and inhibited conditions displayed notably different occurrence distributions (Figures 4D-E), the biggest deviations of which were restricted to specific regions of behavior space (Figure 4F). Control larvae were more likely to visit parts of the behavior space that correspond to pauses and bends (Figure 4F; Figure S3) while inhibited larvae spent more time, proportionally, in the crawling oscillator (Figure 4F). To confirm these differences using a TREBLE independent approach, we compared the primary behavioral features themselves, as measured using the FIM system, and observed that inhibited larvae bend less, crawl further, and have a significantly increased velocity distribution relative to control animals (Figure 4G).

Finally, to quantitatively compare control and transiently inhibited larvae in the TREBLE space, we clustered the behavior space based on similarity to identify discrete elements of behavior that together represent foraging (see Methods). We then examined whether control and transiently inhibited larvae displayed changes in the frequency of occurrence of discrete behavioral motifs (Figure 4H). Reflecting our previous findings, the mechano-ch silenced and control animals displayed different overall distributions (Figure 4I) and significantly varied across a number of behavioral motifs (Figure 4J; trial-wise Kruskal-Wallis test). Specifically, controls were more likely to pause (Figure 4J; dark blue cluster; p < 0.01) and head cast during crawling (Figure 4J; dark orange cluster; p < 0.002) while the mechano-ch silenced larvae were more likely to be crawling (Figure 4J; red, orange, light green clusters). These findings demonstrate that developmental inhibition of mechano-ch neurons leads to an apparent simplification of larval foraging behavior, biasing animals toward ongoing crawling as opposed to the typical sequence of crawling, pausing, and head casting.

## Discussion

These studies demonstrate that PaSNA in the *Drosophila* embryo follows a stereotyped sequence of wave-like, large-scale network activation events interspersed by low-activity periods (Figure 1, 2). Strikingly, our data also demonstrate that muscle contractions shape the magnitude of PaSNA via mechanosensory input, beginning during the myogenic phase, but perhaps also continuing throughout each subsequent wave (Figure 3). Transiently disrupting this mechanosensory feedback during embryonic development results in deficits in larval locomotor behavior, arguing that neural activity plays a critical role in the functional organization of locomotor circuits (Figure 4). These results suggest that sensory inputs generated by spontaneous muscle contraction play a role in subsequent circuit establishment, thereby providing one of the earliest examples of sensory regulation of locomotor development in any context.

Our work has measured the trajectory of neural activity across embryonic development. Prior to PaSNA, individual CNS neurons display transient elevations in intracellular calcium levels (flickering) that depend on depolarization of the plasma membrane, but which are independent of synaptic input. We hypothesize that these cells are spontaneously excitable. In addition, as previous work has demonstrated, muscle twitching independent of neural activity also occurs, and based on our work, appears to lead to the selective activation of mechano-ch sensory neurons. The output of mechano-ch neurons then acts to negatively modulate the basal levels of intracellular calcium in the CNS, the onset of PaSNA, and the amplitude of PaSNA waves. While it is possible that the activation of mechano-ch neurons occurs independently of muscle contraction, given the phenotypic similarities revealed by muscle inhibition and mechano-ch inhibition (Figure 3), we favor a model where mechano-ch neuronal activity is driven by muscle contraction.

Our data demonstrate that the first episode of PaSNA invariably begins in the thoracic region (Figure 2). After this initial event, PaSNA proceeds through a stereotyped sequence of accelerating, intensifying waves through to hatching. Given the striking similarity in both the spatial and temporal properties of PaSNA in the embryo, as well as analogous observations in the *Drosophila* visual system ^21^, we hypothesize that this process is under tight genetic control. Similarly, observations across systems, including in the mammalian cortex, have led to speculation that genetic information underlies the spontaneous neuronal activity present in these developing circuits ^22^. Intriguingly, our observations parallel previous results in the developing chick spinal cord, where waves of activity are also preceded by sporadic activity and PaSNA initiates in a localized region at the anterior part of the spinal cord ^23^. We speculate that evolutionarily ancient mechanisms initiate PaSNA in motor systems in both invertebrates and vertebrates. Our characterization of the initiation and progression of PaSNA in the *Drosophila* embryo sets the stage for the dissection of these mechanisms at the level of specific circuits, cells, and molecules.

Our quantitative measurements of PaSNA put constraints on the molecular basis of its implementation. In particular, the speed with which a single wave of activity traverses the nervous system is remarkably slow, taking approximately 75 seconds to move from the initiation zone to the most posterior region of the VNC. By comparison, the wave of neural activity needed to produce a wave of crawling takes approximately one second to travel the same distance ^24,25^. We infer that wave propagation during embryogenesis is unlikely to proceed by a simple neuron-to-neuron sequence of synaptic transmission events. Understanding the cellular and molecular basis of this wave propagation mechanism represents an important challenge for future work.

Most previous studies examining PaSNA function in shaping developing circuits rely on perturbations that abolish PaSNA. Interestingly, our study shows that an increase in PaSNA also leads to changes in behavior. This result suggests that organisms must quantitatively tune the level of PaSNA during circuit establishment. Supporting this, previous studies have demonstrated that excessive neuronal activity during larval circuit formation leads to hyperexcitable motor circuits that are prone to seizures ^26,27^. It is possible that the excessive PaSNA experienced after mechano-ch inhibition leads to hyperexcitability of specific neurons within the circuits that control foraging behavior. Identifying the neurons affected upon mechano-ch transient inhibition and probing their electrophysiological properties will test this idea.

Relatively little attention has been paid to examining the role of spontaneous neural activity in shaping innate behaviors. In that light, our finding that the activity of mechano-ch neurons during development shapes locomotor behavior is remarkable. Given that this behavioral effect is developmentally programmed, we hypothesize that mechano-ch input is needed to pattern connectivity or determine the physiological properties of specific cells in developing motor circuits. Indeed, blocking synaptic transmission in mechano-ch neurons throughout development changes the connectivity of these cells with their post-synaptic partners, cells that mediate behavioral responses to vibration ^28^. We hypothesize that the changes that mechano-ch inputs exert on developing circuits are, in fact, widespread, modifying circuits across the CNS through PaSNA. Supporting this idea, the locomotor phenotype of inhibiting mechano-ch neurons after PaSNA is very different from our targeted developmental inhibition of the same neurons ^12–14^. In vertebrates, motor feedback is crucial to shaping learned motor behaviors through activity-dependent mechanisms ^29^. It is tempting to speculate that the sculpting of innate foraging behavior by mechano-ch neuron activity in *Drosophila* reveals an analogous, evolutionarily ancient mechanism that may have been co-opted in other contexts to enable motor learning.

## Materials and Methods

### Fly Stocks

All stocks were kept at 25°C on molasses-based food. The following stocks were used: UAS-IVS-Syn21-GCaMP6s-P2A-nls-tdTomato-p10 on JK66B was a gift from Marta Zlatic (MRC Laboratory of Molecular Biology). LexAop-Kir_2.1_ at VIE-260B, UAS-Kir_2.1_ at VIE-260B and LexAop-TNT on VIE-260B were gifts from Barry Dickson (The University of Queensland). pBDP-LexA:p65 on attp40 was a gift from T. Shirangi (Villanova University). UAS-GtACR1 at attP2 was a gift from A. Claridge-Chang (Duke-NUS Med School). The following stocks were obtained from the Bloomington Drosophila Stock Center: elav-GAL4.L on 3rd (BDSC# 8760), elav-GAL4.L on 2nd (BDSC# 8765), elav-GAL4.L on 3rd (BDSC# 8760), GMR44H10-lexa::p65 on attP40 (BDSC# 61543), elav^c155^-GAL4 (BDSC# 458), UAS-TeTxLC.tnt G2 (BDSC# 28838), UAS-mLexA-VP16-NFAT, LexAop-rCD2-GFP (CaLexA) (BDSC# 66542), LexAop-CD8-GFP-2A-CD8-GFP on 2nd (BDSC# 66545), iav-lexA::p65 ^30^ on VK00013 (BDSC# 52246), attP-9A VK000013 (BDSC# 9732) and iav-GAL4.K on 3rd ( BDSC# 52273).

### Embryo collection for calcium imaging

For all imaging experiments, embryos were collected in 15-30 minute time windows the day before imaging, and grown at 25°C or 23°C on standard 3.0% agar molasses collection caps covered with a thin layer of wet yeast. Before imaging, embryos were dechorionated with double-sided tape and staged using elongation of the anterior midgut as a guide ^4,7^. To prevent dehydration, embryos were transferred into Halocarbon oil or saline no more than 5 minutes after dechorionation.

### Wide-field imaging

Staged, dechorionated embryos were mounted ventral side up on double-sided tape, covered with Halocarbon oil (180 cSt) and imaged using a Leica M205 FA system with a Plan Apo Corr. 2X objective. For experiments shown in Figure 3 D-F, embryos were mounted on Sylgard covered with an oxygenated saline solution (103 mM NaCl, 3 mM KCl, 5 mM TES, 1 mM NaH2PO4, 4 mM MgCl2, 1.5 mM CaCl2, 10 mM trehalose, 10 mM glucose, 7 mM sucrose, and 26 mM NaHCO3). Stereoscopic magnification was used to achieve a final magnification of 64X (Figure 1) and 80X (Figure 3 D-F,). Fluorescent signals were acquired using LED illumination (CooLED pE-300 white). GCaMP6s was excited and collected using an ET470/40x ET525/50m band-pass filter set, while tdTomato was excited and collected using an ET545/25x ET605/70m band-pass filter set, acquiring each signal sequentially. Each cycle of imaging acquisition was 7 seconds long. We used a back-thinned sCMOS camera (Orca-Fusion BT - Hamamatsu) to capture images at a 1024 × 1024 resolution (after 2×2 binning), corresponding to a pixel size of 2.0 μm × 2.0 μm (Figure1); and 512 × 512 resolution (after 4×4 binning), corresponding to a pixel size of 3.3 μm × 2.3 μm (Figure 3 D-F). Imaging sessions were from 2 hrs to 9 hrs in duration, depending on the experiment, and were conducted at 23±3°C.

### Two-photon imaging

Staged, dechorionated embryos were mounted ventral side up on Sylgard pads and imaged using a Bruker Ultima system. We used a Leica 20X HCX APO 1.0 NA water immersion objective lens, a piezo objective mount, resonant scanning and GaAsP PMTs. GCaMP6s and tdTomato signals were excited with a Chameleon Vision II laser (Coherent) at 920nm, and collected through a 525/50nm or a 595/50nm filter, respectively. Both signals were simultaneously collected using resonant scanning mode. Imaging volumes were acquired at an XY resolution of 358 × 148 (corresponding to a pixel size of 1.05 μm × 1.05 μm), with 41 z-sections separated by 1.5μm steps, at a volume rate of 2.6Hz. During the entire imaging session embryos were submerged in an oxygenated saline solution (as above), and kept at 25C°.

### Immunostaining and confocal imaging

Immunostaining was performed as previously described ^31^. The 1° antibody used was chicken anti-GFP (1:2,000, Abcam). The 2° antibody used was anti-chicken Alexa 488 (1:500, Life Technologies). Confocal image stacks were acquired on a Leica SP8, using 40X HC PL APO 40X 1.3NA oil objective and a HyD detector. Images were processed in Fiji (https://imagej.net/Fiji). Adjustments to brightness and contrast were applied uniformly to the entire image.

### Behavior data collection

Parents were crossed and fed with wet yeast containing 0.5 mM all *trans*-Retinal (ATR) at least three days before embryo collection. ATR and yeast were replaced every day. Embryos were collected for 30 minutes on standard 3.0% agar molasses collection caps covered with a thin layer of wet yeast without ATR and incubated at 25°C in darkness. 15.5 hours later, embryos were placed under a 3.8uW/mm^2^ 550nm LED for 5 hours. Light pulses 600ms long were delivered at one second intervals, as previously shown to induce inhibition in *Drosophila* embryos using halorhodopsin ^26^. Halorhodopsin, like GtACR1, is a silencing optogenetic tool that relies on chloride ions entering the cell. Control animals were kept in the same incubator, in darkness. One day after light exposure was terminated, at the L1 stage, animals were collected and transferred to a Petri dish with1.0% agar and relocated to a room kept at 23°C and 60% humidity. After 10 minutes of acclimation to the room, groups of 8 to 12 larvae were transferred to a 7.5 x 7.5 cm 1.0% agar arena. After 15 to 30 seconds, locomotion was recorded using a FIM imaging system (^17^ https://www.unimuenster.de) at 10 fps for 5 minutes. The FIM system was equipped with an azA2040-25gm camera (Basler) and a TEC-V7X macro zoom lens (Computar). Individual larvae were then tracked using FIMtrack software ^17,32^. Primary measurements from FIMtrack were used for behavioral analyses (see below).

## Quantitative and statistical analysis

### Processing of calcium imaging data

After image acquisition, regions of interest (ROIs) were manually drawn on the ventral nerve cords and mean intensities were extracted using LAS X software (Leica Microsystems). For Figure 2 ROIs were drawn on Fiji (https://imagej.net/Fiji). To account for movement of the embryo and changes in gene expression over time, we encoded and recorded a structural fluorescent marker (tdTomato) in conjunction with the calcium sensor (GCaMP6s) and considered the ratio of the latter to the former as our measurement of calcium levels in the embryo. This ratiometric calcium signal was then converted into ΔF/F signal, dependent on a baseline signal computed separately for each embryo. For figures 2 and 3, the initial baseline for ΔF/F prior to peak detection was determined by calculating the mean of the 100 values lowest ratiometric values. For figure 1, the baseline for each time point in the ratiometric calcium signal was computed as a function of 16 minutes of the signal flanking the time point of interest (8 minutes prior to and 8 minutes after the time point). The 16-minute signal was divided into 20 bins of signal amplitude ranges. The bin with the largest number of samples was taken to primarily reflect the baseline, while other bins were taken to reflect deviations from the baseline. The choice of 20 bins was made empirically based on the sparsity of neuronal activity. The mean of the samples in the largest bin was considered the baseline value for the time point in the middle. At the two edges of the signal, where the full 8 minutes prior to or after the considered time point do not exist, linear fits were used as the baseline. A 150-second, quartic Savitzky–Golay filter was applied to the resulting ΔF/F signal.

### Episode and peak detection

For figures 2, a 150-second, cubic Savitzky–Golay filter was first applied to the initial ΔF/F trace. Standard deviation for the filtered data was then calculated. Candidate first episodes were detected by finding the first instance where the filtered signal is equal to or greater than 1.2 times the standard deviation. Given that the intervals between episodes are at least 25 minutes, the large increase in signal must appear after a minimum of 30 minutes in order to be considered as a bona fide first episode. These candidate episodes were then manually curated for miss-called episodes due to small fluctuations that resulted in rapid increase in signal but were not sustained over longer than 20 seconds. Traces were then trimmed from −245 to 800 seconds (time series plots) relative to the initiation of the episode. These traces were used to calculate a new ΔF/F with a new baseline that was calculated as the mean of the 25 timepoints with the lowest signal. The Seaborn library was then used to plot traces.

For Figures 1 and 3, where episodes throughout PaSNA were monitored, peaks in the ΔF/F signal were detected using thresholds in the zeroth, first, and second derivatives of the signal. Each derivative signal was filtered with a 150-second, quartic Savitzky-Golay filter. The first derivative threshold was used to detect a rapid rise, while the second derivative was used to detect concavity. First, values crossing the zeroth derivative threshold were identified as peak candidates. A minimum peak distance of 500 seconds was enforced, following a greedy heuristic that kept candidates with the largest values first. Manually analyzed data showed that distinct episodes are invariably separated by more than 600 seconds. Then, of the remaining candidates, only those preceded by threshold crossings in both the first derivative and the second derivative within 210 seconds were kept and detected as episodes. Minimum thresholds of 0.06 for the zeroth derivative and 0.006 for the first derivative were derived empirically, while the maximum threshold of 0 for the second derivative was chosen to select for concavity. For Figure 3 and FigureS2 we calculated the area under the curve 20 seconds surrounding the peak using the trapezoidal rule.

### Stereotypy of episode timing

We assessed the extent to which PaSNA episodes occurred with stereotyped interpeak intervals by comparing the episode interval distributions of embryos to those generated under a null model. Under the null model, episodes corresponding to each real embryo occurred randomly following a Poisson process with a rate parameter equal to the mean rate of the first twelve episodes in the real embryo. In our Monte Carlo sampling of interpeak intervals, we rejected those under 500 seconds, consistent with the minimum peak distance imposed in our peak detection algorithm. We sampled 1,000,000 model embryos for each of 23 real embryos such that more than 500,000 remained after the rejections. We then computed the root mean squared error (RMSE) from the mean for each model dataset of 23 embryos, and the resulting distribution was compared against the RMSE computed for each peak in the real dataset. We assessed significance by examining how many model datasets had RMSE lower than that of each peak in the real dataset, and corrected for multiple comparisons using the Holm-Bonferroni method.

### Statistical analysis

Statistical tests for Figure 3F, I, J, M; Figure S2A, B, E, H, I, J were done with Graphpad Prism. Statistical tests for Figure 3H, L; FIgureS2C, D, G, K were done with scipy and statsmodels libraries. For two group comparisons with equal variance, we conducted unpaired-student’s t-test. For two group comparisons with unequal variance, we conducted Welch’s t-test. For three groups comparisons with unequal variance, we conducted Brown-Forsythe and Welch ANOVA followed by a Dunnett’s T3 multiple comparison test. For multiple comparison of two groups we used an unpaired student’s t-test with Holm-Bonferroni method.

### Behavioral analysis

Primary measurements from FIMtrack ^33^ reflecting larval size, shape, and velocity were used for input, in addition to the angular velocity of the head, midpoint, and tail. Size measurements (i.e. area, perimeter, radii, spine length) were detrended using the ma function in the R package forecast (window size = 10) and converted to z-scores. Principal component analysis was used to control for potentially redundant information in the input features, yielding 8 principal components that explained >90% of the variance in the feature set. To find the appropriate timescale with which to analyze the behavioral features an empirical window search procedure was used (described in ^18^). We constructed behavior spaces using the top 8 PCs sweeping window sizes ranging between 100 ms and 5 seconds. For a given window size (denoted w), the windows were compiled as follows: given frame i, the 8 PCs corresponding to frames i:i+w were linearized and concatenated, resulting in a vector with length 8w. This was repeated for all windows in the data set and the resulting vectors were appended to produce a window matrix with 8w rows. A behavior space was then constructed by embedding this matrix into low-dimensional space via the UMAP algorithm ^34^. The appropriate window size was then determined by comparing the structural (Procrustes and Euclidean distance) and temporal features (recurrence) of behavior spaces produced from 20 random trials per window size. As was found before ^18^, a window size of 800ms was chosen.

We then created a behavior space encompassing the full control and transient inhibition datasets using this window size. Trials were first filtered to include those that were longer than 2.5 seconds and that traveled at least 50mm, resulting in 84 control and 97 transient inhibition trials and a total of 179,409 frames. The resulting behavior space captured the major components of the larval foraging ethogram (Figures 4B-C). Differences in behavior patterns between the conditions were inferred using 2-dimensional kernel density estimation (as in Figures 4B-C) computed over all trials for each condition. The difference map in Figure 4F was produced by first normalizing via division by the greatest value (to produce a range of values between 0 and 1) and then subtracting the transient inhibition map from the control map. Differences in individual feature distributions (as in Figure 4G) were assessed using a Kruskal-Wallis test comparing the mean value for each trial across conditions (for each test n = 84 control and n = 97 for inhibited). To control for the autocorrelation in behavior we sampled each measurement every 10th frame for a total of ~ 8,000 measurements per condition.

Louvain clustering was used to identify discrete components of behavior space. First, a graph was created with 2 sets of edges: the first representing the xy-coordinates in behavior space for each frame and the second corresponding to the xy-coordinates of the immediately following frame. This graph provides both information about the local neighborhood densities of the points in behavior space and the temporal sequencing between points over time. Louvain clustering was then run on this graph using the function cluster_louvain in the R package igraph ^35^. Differences in occurrence in each cluster between the conditions were assessed using a Kruskal-Wallis test, again comparing the occurrence density of all individual trials between the two conditions (n = 84 control and n = 97 for inhibited).

## Acknowledgements

We thank D. Berfin Azizoglu for comments on the manuscript. Stocks obtained from the Bloomington Drosophila Stock Center (NIH P40OD018537) were used in this study. Funding was provided by HHMI (CQD, AC-R), NIH HD27056 (CQD), F32NS105350-01A1, The Walter V. and Idun Berry Postdoctoral Fellowship, K99NS119295-01 (AC-R), the Stanford School of Medicine Dean’s Fellowship (RAY), The Simons Foundation (TRC), and the National Defense Science & Engineering Graduate Fellowship, Stanford Graduate Fellowship, and Stanford Mind, Brain, Computation and Technology Training Program (MC).

## Author contributions

Conceptualization, AC-R, CQD and TRC; Methodology: AC-R; Software, AC-R, RAY and MC; Formal analysis, AC-R, RAY and MC; Investigation, AC-R; Writing, AC-R, CQD and TRC; Visualization, AC-R, RAY and MC

---

**Figure S1 related to Figure 1.**
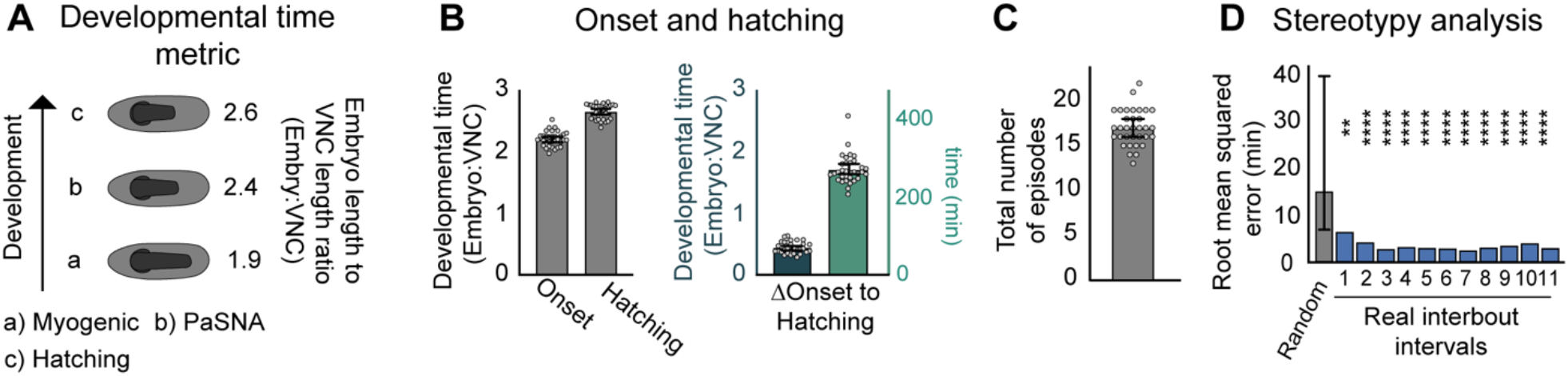
Onset and length of PaSNA quantification. **(A)** Developmental time metric used to quantify PaSNA onset and progression. **(B)** Onset and hatching measurements of PaSNA (n = 33). **(C)** Number of total episodes from PaSNA onset to hatching (n = 33). Bar plots represent mean with 95% confidence interval. **(D)** Interval stereotypy analysis in which the distribution of interpeak intervals across all embryos are compared to Poisson processes with the same mean (see Methods). ** p < 0.01, *** p< 0.001, **** p < 0.0001; two-sample t-test, Holm-Bonferroni correction.

**Figure S2 related to Figure 3.**
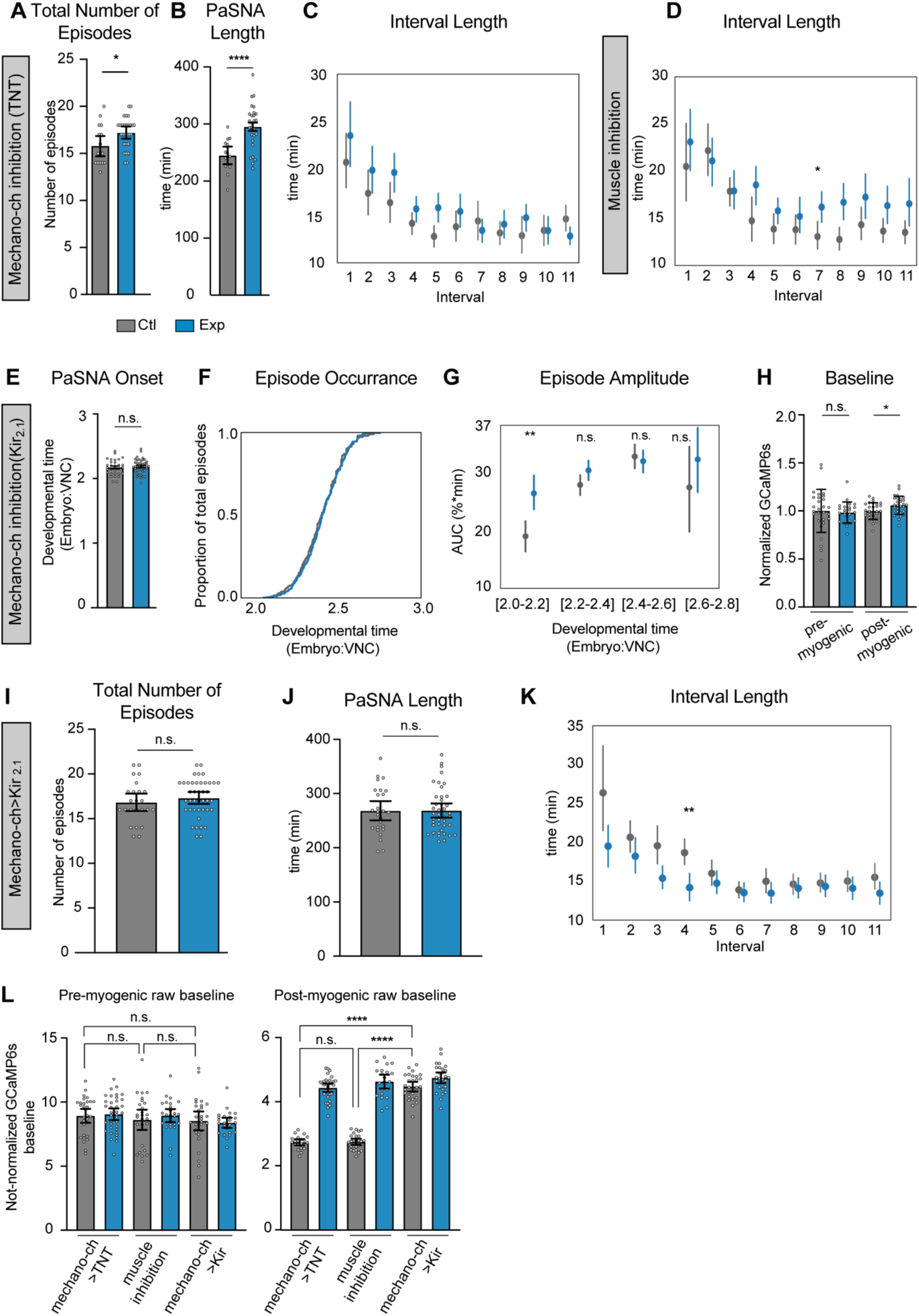
Mechanosensory neurons modulate the amplitude of PaSNA episodes. **(A-C)** Quantification of PaSNA phenotypes in control embryos in gray and experimental embryos expressing TNT in mechano-ch in blue. **(A)** Total number of episodes from PaSNA onset to hatching (n = 16 control; n = 28 experimental). **(B)** Time span from PaSNA onset to hatching (n =16 control; 28 experimental). **(C)** Quantification of the first eleven interbout interval length (n = 17 control; n = 32 experimental). **(D)** Quantification of the first eleven interbout interval lengths (n = 28 control; n = 28 experimental) for control embryos in gray and experimental embryos expressing Kir_2.1_ in muscles in blue. **(E-K)** Measurements of the timing and intensity of PaSNA for control embryos (gray) and experimental embryos expressing Kir_2.1_ in mechano-ch neurons (blue). **(E)** Quantification of PaSNA onset (n = 30 control; n = 41 experimental). **(F)** Cumulative occurrence of the first twelve episodes plotted as the proportion of total episodes across developmental time (n = 27 control; n = 29 experimental). **(G)** AUC quantification for the first twelve episodes plotted against binned developmental time (n = 26, control; n = 41 experimental). **(H)** Quantification of GCaMP6s baseline levels normalized against control mean before (n = 28 control; n = 24 experimental) and after the myogenic phase (n = 28 control; n = 28 experimental). **(I)** Total number of episodes from PaSNA onset to hatching (n = 24 control; n = 41 experimental). **(J)** Time span from PaSNA onset to hatching (n =26 control; n = 41 experimental). **(K)** Quantification of the first eleven interbout interval lengths (n = 27 control; n = 29 experimental). **(L)** Raw GCaMP6s baselines for control (gray) and experimental (blue) groups. Different experiments labeled on the X axis. As GCaMP6s and TdTomato are expressed at low levels during the pre-myogenic stage, we increased excitation power at this stage, making direct comparisons between pre- and post-myogenic stages impossible. For all point plots, points represent mean and lines depict the 95% confidence interval. For all bar graphs the mean and 95% CI are displayed. ****p<0.0001, ***p<0.005, **p<0.005, *p<0.05. For (**A**), (**B**), (**E**), (**I**) and (**J**) we used a two-sample t-test. For (**C**), (**D**), (**G**) and (**K**) we used two-sample t-tests with Holm-Bonferroni correction. For (**H**) we used two-sample Welch’s t-tests to account for difference in variance. For (**L**) we used a Brown-Forsythe and Welch ANOVA followed by a Dunnett’s T3 multiple comparison test to account for differences in variance. For genotypes information see Table S1.

**Figure S3 related to Figure 4.**
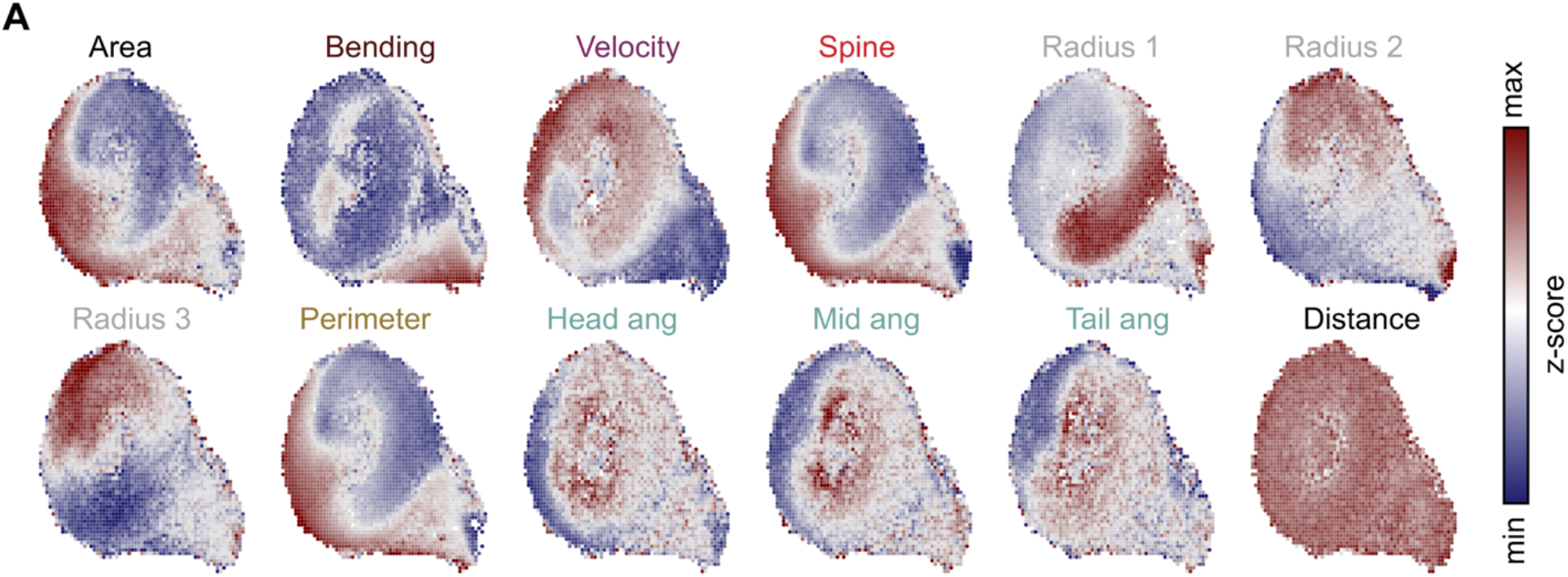
Primary behavioral metrics as a function of larval behavioral space. The distribution of area, bending, velocity, spine length, radius 1, radius 2, radius 3, perimeter, head angle, middle body angle, tail angle, and distance as a function of larval behavior space (z-scores). Z-score scale to the right.

**Video S1 High-throughput wide-field calcium imaging.** Pan-neuronal GCaMP6s (top) and TdTomato (bottom) signals of an individual representative embryo throughout PaSNA at 750 times real time speed. Video is an XY cropped region from the time-lapse used for the snapshot shown on Figure 1. Scale bar is 100μm and the timestamp is in minutes:seconds.

**Video S2 Two-photon calcium imaging of the first PaSNA episode.** Maximum intensity projections of pan-neuronal GCaMP6s signals in a representative control embryo imaged using two photon microscopy.

**Video S3 PaSNA depends on depolarizations.** Maximum intensity projections of pan-neuronal GCaMP6s signals in a representative embryo expressing Kir2.1 pan-neuronally, imaged using two photon microscopy.

**Video S4 PaSNA depends on synaptic transmission.** Maximum intensity projections of pan-neuronal GCaMP6s signals in a representative embryo expressing TNT pan-neuronally, imaged using two photon microscopy.

